# Extended Application of Genomic Selection to Screen Multi-Omics Data for Prognostic Signatures of Prostate Cancer

**DOI:** 10.1101/2020.06.02.115816

**Authors:** Ruidong Li, Shibo Wang, Yanru Cui, Han Qu, John M. Chater, Le Zhang, Julong Wei, Meiyue Wang, Yang Xu, Lei Yu, Jianming Lu, Yuanfa Feng, Rui Zhou, Yuhan Huang, Renyuan Ma, Jianguo Zhu, Weide Zhong, Zhenyu Jia

## Abstract

Prognostic tests using expression profiles of several dozen genes help provide treatment choices for prostate cancer (PCa). However, these tests require improvement to meet the clinical need for resolving overtreatment which continues to be a pervasive problem in PCa management. Genomic selection (GS) methodology, which utilizes whole-genome markers to predict agronomic traits, was adopted in this study for PCa prognosis. We leveraged The Cancer Genome Atlas (TCGA) database to evaluate the prediction performance of six GS methods and seven omics data combinations, which showed that the Best Linear Unbiased Prediction (BLUP) model outperformed the other methods regarding predictability and computational efficiency. Leveraging the BLUP-HAT method, an accelerated version of BLUP, we demonstrated that using expression data of a large number of disease-relevant genes and with an integration of other omics data (*i.e*., miRNAs) significantly increased outcome predictability when compared with panels consisting of small numbers of genes. Finally, we developed a novel stepwise forward selection BLUP-HAT method to facilitate searching multi-omics data for predictor variables with prognostic potential. The new method was applied to the TCGA data to derive mRNA and miRNA expression signatures for predicting relapse-free survival of PCa, which were validated in six independent cohorts. This is a transdisciplinary adoption of the highly efficient BLUP-HAT method and its derived algorithms to analyze multi-omics data for PCa prognosis. The results demonstrated the efficacy and robustness of the new methodology in developing prognostic models in PCa, suggesting a potential utility in managing other types of cancer.

## Introduction

Prostate cancer (PCa) is the second most common cancer in men worldwide. An estimated 1,276,106 new cases and 358,989 deaths were reported in 2018 [1]. Three major challenges need to be better addressed through biomarker studies to improve the management of the disease and save lives: (I) early detection of the disease, (II) accurate prediction of tumor progression to avoid overtreatment, and (III) guidance for personalized therapies for patients carrying different subtypes of PCa. With a focus on the second challenge, this study adopted the methodology of genomic selection/prediction (GS), which is commonly applied in agricultural breeding, for an integration of multi-omics to improve the predictive ability (or predictability, defined in the Methods) for PCa prognosis.

The majority of PCa tumors grow slowly and will likely never cause health problems. A small percentage of patients carry aggressive PCa and require immediate treatment. Patients with slow growing tumors only require active surveillance. Lacking effective tests to provide patients with the best choices for treatment based on their individual disease states, overtreatment continues to be a health issue in PCa management owing to the associated negative and unnecessary side effects. A few clinically applicable gene expression signatures have been developed to calculate risk scores for PCa prognosis, including Prolaris (Myriad Genetics Inc.), a gene expression signature assay that is based on 31 genes involved in cell cycle progression for cancer risk stratification [2], Decipher (GenomeDx Biosciences Inc.), a 22-marker expression panel for prediction of systemic progression after biochemical recurrence [3], and OncotypeDX Genomic Prostate Score (Genomic Health, Inc.,), which consists of 17 genes (12 selected genes in four biological pathways and five reference genes) to predict adverse pathology at the time of radical prostatectomy [4]. Compared with the clinically applied nomograms [5], these multiple-gene tests only provide a moderate improvement to disease prognosis, and they all need further validation by prospective trials [6, 7]. This leaves a wide gap between clinical practice and its objective for eliminating unnecessary surgeries.

Many common human diseases, including cancer, have a polygenic nature, *i.e*., the disease phenotypes are controlled by many genetic variants with minor effects. Numerous studies have indicated that using genome-wide markers as predictors yielded much higher predictability of complex traits than using a few major Quantitative Trait Loci (QTLs) only [7–11]. The mediocre predictive abilities of the current prognostic tests are likely due to the limited number of genes being included in simple linear models, even though some of these genes are major players of cancer progression. Conventional statistical methods usually cannot efficiently handle highly saturated models with *p » n*, where *p* is the number of parameters (selected markers) of the models and *n* is the sample size. GS is a powerful tool in the fields of plant and animal breeding, which estimate genetic effects of thousands of genome-wide markers simultaneously using whole-genome regression (WGR) models [12, 13]. Numerous advanced statistical methods, including BLUP [14, 15] and Bayesian models (*i.e*., BayesA, BayesB, and BayesC, etc.) [12, 13, 16, 17] have been proposed [18, 19], and the vast success of GS in plant and animal sciences gave an impetus to introduce this powerful application to human medicine.

In this study, we established a novel method, named Stepwise Forward Selection using BLUP-HAT (SFS-BLUPH), and applied this method to data from the TCGA Prostate Adenocarcinoma (TCGA-PRAD) project to develop a multi-omics signature for PCa prognosis. At first, the pre-radical prostatectomy nomogram developed by Memorial Sloan Kettering Cancer Center (MSKCC) was used to derive six quantitative disease traits, including progression-free probability in five years (PFR5YR), progression-free probability in ten years (PFR10YR), organ-confined disease (OCD), extracapsular extension (ECE), lymph node involvement (LNI), and seminal vesicle invasion (SVI). These six traits were then used to evaluate six GS models and three types of omics data including mRNA transcriptome (TR), miRNAs (MI), and methylome (ME) as well as all possible combined data (TR+MI, TR+ME, MI+ME, TR+MI+ME) to identify the best combination of model and omics data for predicting PCa outcomes. The six GS models included BLUP [14, 15], Least Absolute Shrinkage and Selection Operator (LASSO) [20], Partial Least Squares (PLS) [21], BayesB [13], Support Vector Machines (SVM) [22] using the radial basis function (SVM-RBF), and the polynomial kernel function (SVM-POLY). The results indicated that the most widely used GS model, BLUP, outperformed the other models in terms of predictability and computational efficiency. The computational efficiency was further boosted by adopting the BLUP-HAT method, an optimized version of BLUP [23]. With the BLUP-HAT method and the TCGA-PRAD data, we demonstrated that: (I) prediction models using expression profiles of a large number of genes selected from the transcriptome outperformed three clinically employed tests which only considered the expression of a small number of major genes. (II) The predictability for disease traits can be further increased if the selective predictors from other omic types (*i.e*., miRNAs in this study) were also factored into the prognostic models. Finally, we utilized the new SFS-BLUPH method to screen the gene and miRNA expression data in the TCGA-PRAD training dataset for the optimal signatures of predictor variables in predicting RFS followed by a rigorous validation in six independent PCa cohorts. The new SFS-BLUPH methodology demonstrated its translational potential and may be widely adopted for management of other types of cancer.

## Methods

### TCGA-PRAD dataset

Multi-omics data (including HTSeq-Counts of RNA-seq, BCGSC miRNA Profiling of miRNA-seq, and Beta value of Illumina Human Methylation 450 array) and clinical data for 495 PCa patients from the TCGA-PRAD project were downloaded and processed by a series of functions in the R package *GDCRNATools* [24]. The mRNAs and miRNAs with counts per million reads (CPM) <1 in more than half of the patients as well as the methylation probes with any missing values were filtered out before subsequent analysis. Certain clinical characteristics, such as pre-operative PSA, which were not available in the Genomic Data Commons (GDC) data portal were retrieved from Broad GDAC Firehose (https://gdac.broadinstitute.org/). The TCGA-PRAD dataset was used for two purposes: (1) to compare the performance of GS models and different omics data in predicting PCa outcomes and evaluate the predictabilities of tens of thousands of BLUP-HAT models with various numbers of genes or miRNAs, and (2) to serve as a training dataset for the development of a multi-omics signature for RFS prediction. The clinical characteristics for 495 patients were summarized in Table 1.

**Table 1:** Clinical characteristics of the patients in TCGA-PRAD project.

| | | Patients ( $N = 495$ ) |
| --- | --- | --- |
| Age at diagnosis (years) | $\leq 65$ | 353 |
| | $> 65$ | 142 |
| Clinical tumor stage | T1a | 1 |
|  | T1b | 2 |
|  | T1c | 172 |
|  | T2a | 54 |
|  | T2b | 54 |
|  | T2c | 50 |
|  | T3a | 36 |
|  | T3b | 17 |
|  | T4 | 2 |
| Gleason score | $\leq 6$ | 45 |
|  | 7 (3+4) | 149 |
|  | 7 (4+3) | 98 |
| | $\geq 8$ | 203 |
| Pre-operative PSA (ng/mL) | 0-3.9 | 52 |
|  | 4-9.9 | 273 |
|  | 10-19.9 | 99 |
| | $\geq 20$ | 55 |

### Independent validation datasets

The profiling data of mRNAs and/or miRNAs as well as clinical data (with available RFS data) in six public datasets (GSE70769, DKFZ2018, GSE116918, GSE107299, GSE54460, and MSKCC2010) were used to validate the prognostic signatures [25–30]. MSKCC2010 had both mRNA and miRNA data, while the other five datasets only had mRNA data. Detailed information for these six datasets was summarized in Table 2.

**Table 2:** Information of the six publicly available independent validation datasets.

| Dataset | Sample Size | Transcriptome Platform | miRNA Platform | Tissue |
| --- | --- | --- | --- | --- |
| GSE70769 | 85 | Illumina HumanHT-12 V4.0 | × | Fresh frozen |
| DKFZ2018 | 32 | Illumina HiSeq 2000 (RNAseq) | × | Fresh frozen |
| GSE116918 | 229 | ADXPCv1a520642 | × | FFPE |
| GSE107299 | 94 | Affymetrix Human Gene 2.0 ST Array | × | Fresh frozen |
| GSE54460 | 90 | Illumina HiSeq 2000 (RNAseq) | × | FFPE |
| MSKCC2010 | 61 (40)* | Affymetrix Human Exon 1.0 ST Array | Agilent-019118<br>Human miRNA<br>Microarray 2.0 | Fresh frozen |

\* For MKSCC2010 dataset, 61 patients have gene expression data, and 40 of them have both gene expression and miRNA expression data.

Processed microarray data for GSE70769 and GSE116918 were downloaded from GEO (https://www.ncbi.nlm.nih.gov/geo/) using R package *GEOquery* [31]; Reads per kilobase per million mapped reads (RPKM) data for DFKZ2018 and processed microarray datasets for MSKCC2010 were downloaded from cBioPortal (https://www.cbioportal.org/) [32]. Raw data of GSE107299 were downloaded from GEO and normalized with the Robust Multichip Average (RMA) method implemented in the R package *oligo* [33]. Raw sequencing data for GSE54460 were downloaded from SRA (https://www.ncbi.nlm.nih.gov/sra) under the accession number SRP036848. The raw sequencing data were aligned using *STAR* (*version 2.7.2a*) software [34], quantified using *featureCounts* (*version 2.0.0*) software [35], and normalized using the Trimmed Mean of M-values (TMM) normalization method implemented in the R package *edgeR* [36].

### Pre-radical prostatectomy nomograms

The pre-radical prostatectomy nomogram (https://www.mskcc.org/nomograms/), developed by the MSKCC, utilizes pre-treatment clinical data to predict the extent of the cancer and long-term outcomes following radical prostatectomy, which can be analyzed as quantitative traits by genomic prediction models. We used this tool to predict six post-surgery disease traits, including progression-free probability in five years (PFR5YR), progression-free probability in ten years (PFR10YR), organ-confined disease (OCD), extracapsular extension (ECE), lymph node involvement (LNI), and seminal vesicle invasion (SVI). The pre-surgery clinical characteristics used for nomogram calculation included age, preoperative PSA level, Gleason score (primary Gleason and secondary Gleason), and clinical tumor stage based on the American Joint Committee on Cancer (AJCC) version 7 staging system [37].

### Genomic selection methodologies

In this study, we compared the predictive ability of six widely used GS methods, including BLUP, LASSO, PLS, BayesB, SVM-POLY, and SVM-RBF. The BLUP method was implemented using a custom R script [38]. LASSO, PLS, and BayesB were implemented in the R packages *glmnet* [39], *pls* [40], and *BGLR* [41], respectively. The two SVM methods, SVM-RBF and SVM-POLY, were implemented in the R *kernlab* package [42].

The mRNA, miRNA, and methylation features, which were initially profiled in different ranges, were rescaled by z-score transformation, allowing for an objective comparison among these multi-omics profiles and for integrated analyses.

The predictability of a model, defined as the squared correlation coefficient (*r^2^*) between the observed and predicted trait values, was calculated through a 10-fold cross validation (CV) procedure. In a 10-fold CV, the sample was arbitrarily partitioned into ten portions with approximately equal size. In each iteration, nine portions were used as the training data to develop the model and the remaining one portion was used as the test data for model evaluation. This process was repeated ten times with each portion having been used as the test data exactly once. The entire 10-fold CV was then replicated ten times to reduce the variation caused by random partitioning.

### BLUP-HAT method

The BLUP-HAT model [23], which produces the same results as BLUP but enjoys much more computational efficiency due to the avoidance of the time-consuming CV, was used in place of the conventional BLUP method to compare the predictabilities of many thousands of models with various numbers of predictors. The linear mixed model that accounts for the relationship between each trait and predictor variables can be expressed as

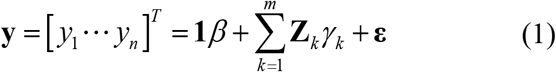

where 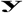 is the vector of trait values for *n* patients, 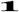 is a vector of 1’s, *β* is the intercept (overall mean), **Z**_*k*_ is a numerical vector for the k^th^ predictor variable, *γ_k_* is the effect of k^th^ variable, *m* is the number of predictor variables in the model, and **ε** is an *n*×1 vector of random errors. We assume that 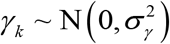 for all *k* = 1,.,.,m, and **ε** ~ **N**(**0, I*σ***^2^) so that

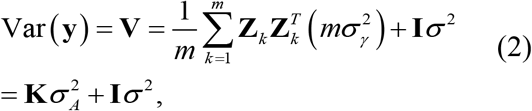

where

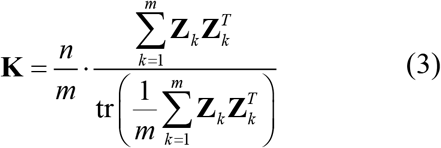

is a relatedness matrix which is equivalent to the kinship matrix in GS [38]. Let us define 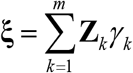 as the poly-predictor effect, and 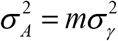 as the poly-predictor variance, we can rewrite the mixed model (1) as

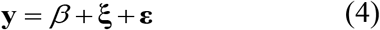

Thence, the Henderson’s equation for the mixed model (4) can be derived as

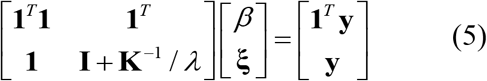

where 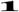 is an identity matrix and 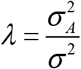. The best linear unbiased estimation (BLUE) of the fixed effects and the best linear unbiased prediction (BLUP) of the random poly-predictor effect are obtained via

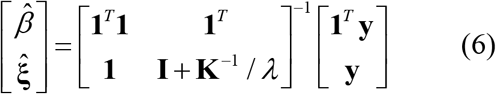

The variance-covariance matrix of the BLUE and BLUP is

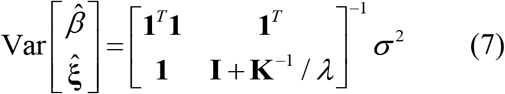

Following the BLUP-HAT method described by Xu [23], the predicted poly predictor effect can be expressed using a linear function of the observed poly-predictor effect involving the hat matrix 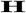, *i.e*.,

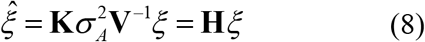

with 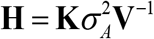. Let 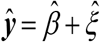 be the predicted trait values and let 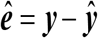 be the residuals, with 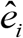 being the *i*^th^ element of the residual vector 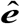. The predicted residual for individual *i* becomes

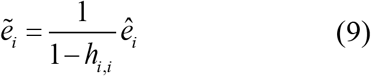

where *h_i,i_* represents the *i*^th^ diagonal entry on 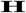. The total sum of squares is defined as

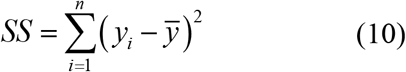

where 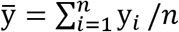.

The predicted sum of squares is

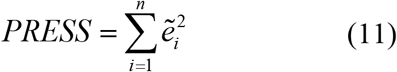

The trait predictability of the BLUP-HAT version is

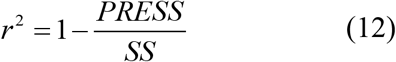

### Commercial panels for PCa prognosis

Three commercial gene expression panels for PCa prognosis were compared in this study, including:

I. Prolaris^®^ (Myriad Genetics Inc., Salt Lake City, US): The Prolaris gene signature consists of 31 cell cycle genes and 15 house-keeping genes. All of the 31 genes can map to Ensembl gene IDs in the TCGA gene expression dataset (Supplementary Table S1). The 15 house-keeping genes were not included in the panel for prediction.
II. Decipher^®^ (GenomeDX Inc., Vancouver, Canada): The Decipher is a 22-marker panel involving 19 genes because two markers may be derived from the same gene (e.g., one in the coding region, and the other one in the intronic region). One of the 19 genes, Prostate Cancer Associated Transcript 32 (PCAT-32) does not have a unique ID in the Ensembl genome annotation, so expression of 18 genes with unique Ensembl IDs were used to represent this panel (Supplementary Table S2).
III. OncotypeDX GPS^®^ (Genomic Health Inc., Redwood City, USA): OncotypeDX GPS consists of 17 genes (12 genes in four biological pathways and five reference genes). Expression of the 12 genes were all quantified in the TCGA dataset and were used for prediction (Supplementary Table S3).

## Results

### Comparison of GS methodologies using various omics data for PCa outcome prediction

We first used the six nomogram-derived traits to systematically evaluate six different GS methods with combinations of various types of omics datasets in full loads (*i.e*., entire mRNA transcriptome, and/or entire set of miRNAs, and/or entire methylome). Although the most important trait of interest for PCa prognosis is the observed clinical outcome (*i.e*., RFS), the nomogram-derived traits can represent collective characteristics of a patient’s disease status and are much less affected by post-surgery therapies compared to the observed outcomes that are sometimes biased and complicated by incorrectly documented treatment history. The MSKCC pre-radical prostatectomy nomogram predicts the extent of the cancer and long-term results following radical prostatectomy, which can be treated as quantitative traits by the GS models. From the TCGA-PRAD dataset, 289 of the 495 primary tumor patients with the available clinical data required for nomogram calculation were used for the analyses. Cox Proportional-Hazards (CoxPH) survival analysis was performed to measure the association between each nomogram-derived trait and RFS. We also performed Kaplan Meier (KM) survival analysis by classifying patients into two risk groups based on the median value for each trait. For PFR5YR, PFR10YR, and OCD, the higher the nomogram values, the lower the risk according to the definitions of the traits. On the contrary, the higher the nomogram values for ECE, LNI, and SVI, the higher the risk. Both CoxPH and KM survival analyses indicated that all the six nomogram-derived traits were significantly associated with RFS (Figure 1), indicating that they were ideal substitutes for the target traits and could be used for evaluating prognostic models.

**Figure 1.**
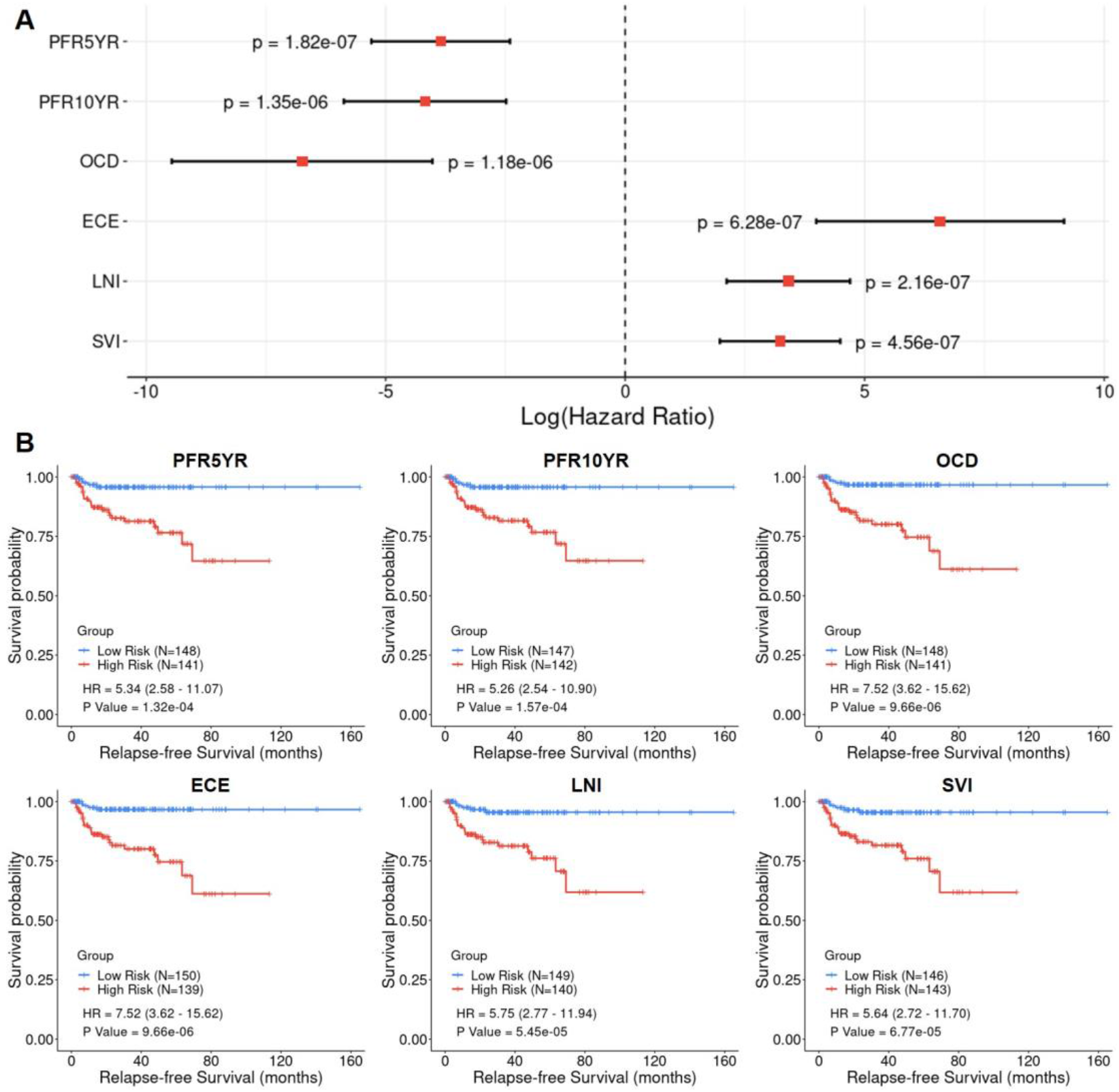
Cox Proportional-Hazards (CoxPH) and Kaplan-Meier (KM) survival analyses of relapse-free survival (RFS) using the six nomogram-derived traits as variables in the TCGA-PRAD dataset. (**A**) Forest plot visualizing the hazard ratio (HR) in log scale, 95% confidence intervals in log scale, and p value of CoxPH survival analysis (**B**) KM curves visualizing the survival probabilities over time for high and low risk groups classified based on the median value of the nomogram-derived scores for each trait.

In total, 285 out of the 289 patients with all the omics data available were used to evaluate the performance of different GS methods and combinations of various types of omics data in predicting nomogram-derived traits. A total of 15,536 genes, 388 mature miRNAs, and 381,602 methylation probes were included for the comparison. The predictabilities of six nomogram-derived traits for the 285 patients were evaluated using six statistical methods and seven omics data combinations *via* 10-fold CV. The results indicated that the predictabilities of different traits varied substantially (Figure 2), with PFR5YR and PFR10YR having the greatest predictabilities. Prediction using mRNA transcriptomic data (TR) outcompeted prediction using either miRNA predictors (MI) or methylome predictors (ME). The combined use of TR and MI in a single model predicted disease outcomes slightly better than the model of using TR alone. In general, prediction models using ME had lower predictabilities than those using TR, MI, and other data combinations. Among the six GS methods, the conventional BLUP method generally outperformed the other methods in terms of trait predictability. In addition, BLUP appeared to be much more efficient in computation time than other methods, especially when a large number of features were included in the models (Table 3). Therefore, the BLUP method as well as the gene and miRNA expression data were selected for the subsequent analyses.

**Figure 2.**
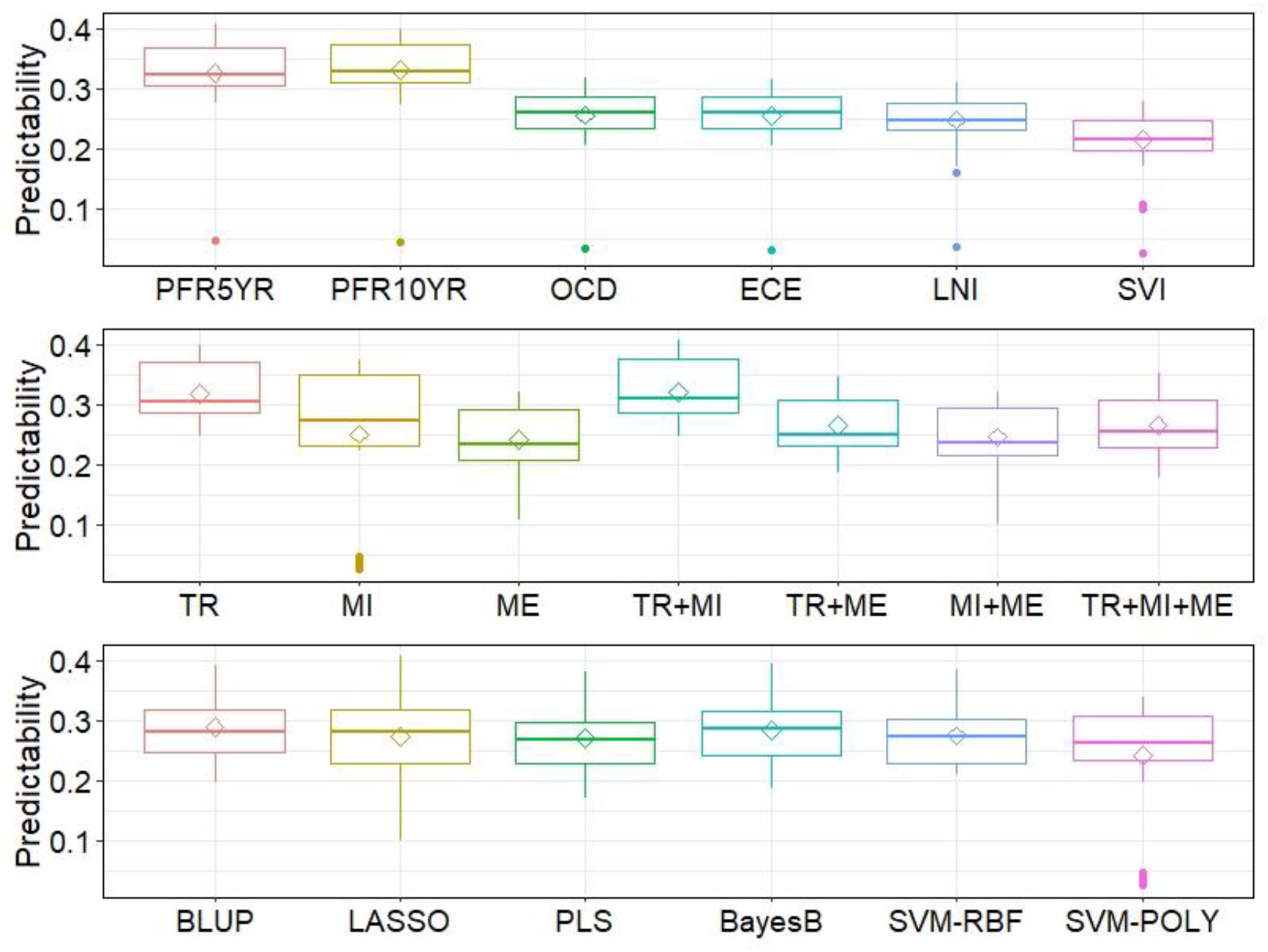
Comprehensive evaluation of the performance of six different genomic selection models (BLUP, LASSO, PLS, BayesB, SVM-POLY, and SVM-RBF) with three omics data (TR: Transcriptome; MI: miRNAs; ME: methylome) and their combinations (TR+MI, TR+ME, MI+ME, and TR+MI+ME) using the six nomogram post-surgery traits (PFR5YR: progression-free probability in 5 years; PFR10YR: progression-free probability in 10 years; OCD: organ-confined disease; ECE: extracapsular extension; LNI: lymph node involvement; SVI: seminal vesicle invasion).

**Table 3.** Computational times in seconds for the six GS models using different omics data (DELL desktop with 16 cores × 2G memory)

| Method | TR | MI | ME |
| --- | --- | --- | --- |
| BLUP | 5 | 1 | 63 |
| LASSO | 34 | 3 | 333 |
| PLS | 104 | 1 | 1,738 |
| BayesB | 385 | 15 | 9,343 |
| SVM-RBF | 145 | 3 | 3,965 |
| SVM-POLY | 149 | 47 | 3,837 |

TR: Transcriptome (15,536 genes); MI: miRNAs (388 mature miRNAs); ME: Methylome (381,602 probes)

### Evaluation of prognostic models with different numbers of genes and/or miRNAs

Enlightened by the report that HAT method yielded the approximate calculation of predictability as the conventional CV in the mixed model analysis but with much improved computational efficiency [23], a BLUP-HAT method was adopted to test tens of thousands of models to test the two proposed hypotheses: (I) using a large number of genes selected from the transcriptome to predict the outcomes of PCa patients will outperform the clinically employed prognostic tests which only rely on several dozen major genes, and (II) the predictive power will be further increased if other omics predictors are also factored into the prognostic models.

The transcriptomic data were used to test the first hypothesis. For each nomogram-derived trait, genes were sorted in descending order based on their absolute Pearson’s correlation coefficients with the trait. Top *N* genes (*N* ranges from 5 to 15,536) selected from the sorted list were sequentially included in the mixed model to calculate the HAT value (predictability, defined in Equation 12 in the Methods section). In each plot of Figure 3, the predictabilities for the models with the top 12, top 18, and top 31 genes, respectively, and the predictabilities for the models consisting of genes in the three commercial tests were marked. We also included a set of control models with 12, 18, and 31 random genes, respectively. For each control model, the random genes were repeatedly selected from the transcriptome ten times, and the average predictability was calculated and labeled by solid lines with different colors in Figure 3. The results indicated that, as expected, the predictabilities of the three commercial panels were significantly higher than the randomly selected genes, confirming the prognostic abilities of those gene panels. It was observed that all the evaluated models with sorted genes being sequentially added had better predictabilities than the three commercial gene panels. The predictabilities rose as more and more genes had been included in the model until they reached the maximum value, where thereafter the predictability values started decreasing. Generally, a few hundred genes were required to have the maximum predictability for each trait, which supported our first hypothesis that the outcome predictability may be substantially boosted by including hundreds of the genes on the top of the sorted gene list when compared with the models using only a small number of the top ‘major’ genes.

**Figure 3.**
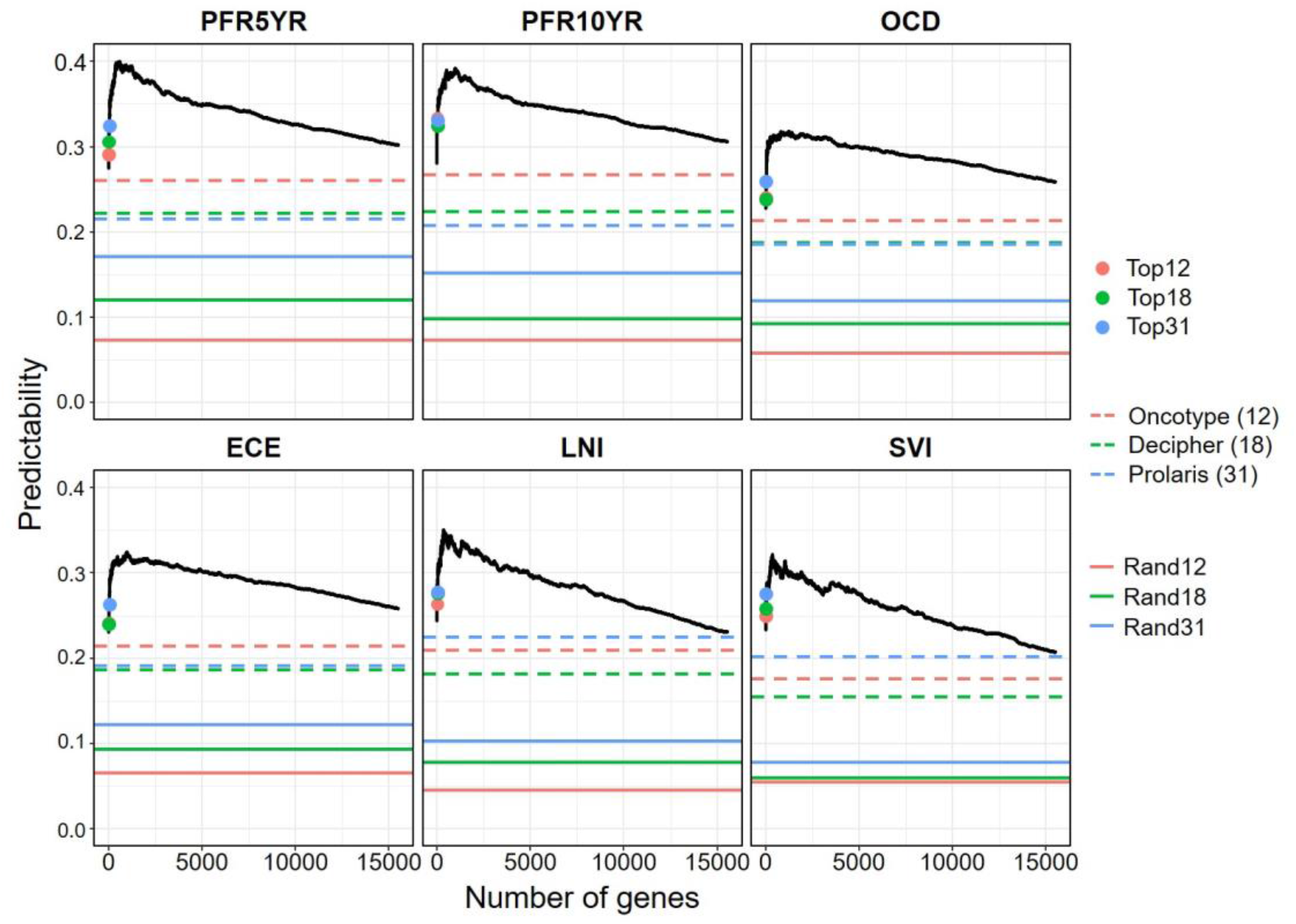
Evaluation of prediction models using different number of genes selected from the transcriptome in predicting six nomogram-derived traits by the BLUP-HAT method. (Top12, Top18, and Top31 represent the top 12, 18, and 31 genes in the ranked gene list, respectively. Rand12, Rand18, and Rand31 represent randomly selected 12, 18, and 31 genes from the transcriptome, respectively). The numbers of genes that achieved the maximum predictabilities for PFR5YR, PFR10YR, OCD, ECE, LNI, and SVI are 470, 995, 1246, 989, 366, and 363, respectively.

To test the second hypothesis that the predictability can be further improved by integrating panels from other omics data, BLUP-HAT was also used to identify the optimal set (top *N*) of miRNAs that reached the maximum predictability. Then the predictabilities of the optimal gene set, the optimal miRNA set, and their combinations were compared for the six traits. The results indicated that: (1) the models using gene expression data outperformed the models using expression data of miRNAs, and (2) the models with combined expression of genes and miRNAs had greater predictabilities than those using genes only, supporting our second hypothesis (Figure 4). To this point, we have used PCa data to successfully provide strong evidence supporting the two hypotheses, which would generally hold in other types of cancers and may help guide the development of improved cancer prognostic models leveraging multi-omics data.

**Figure 4.**
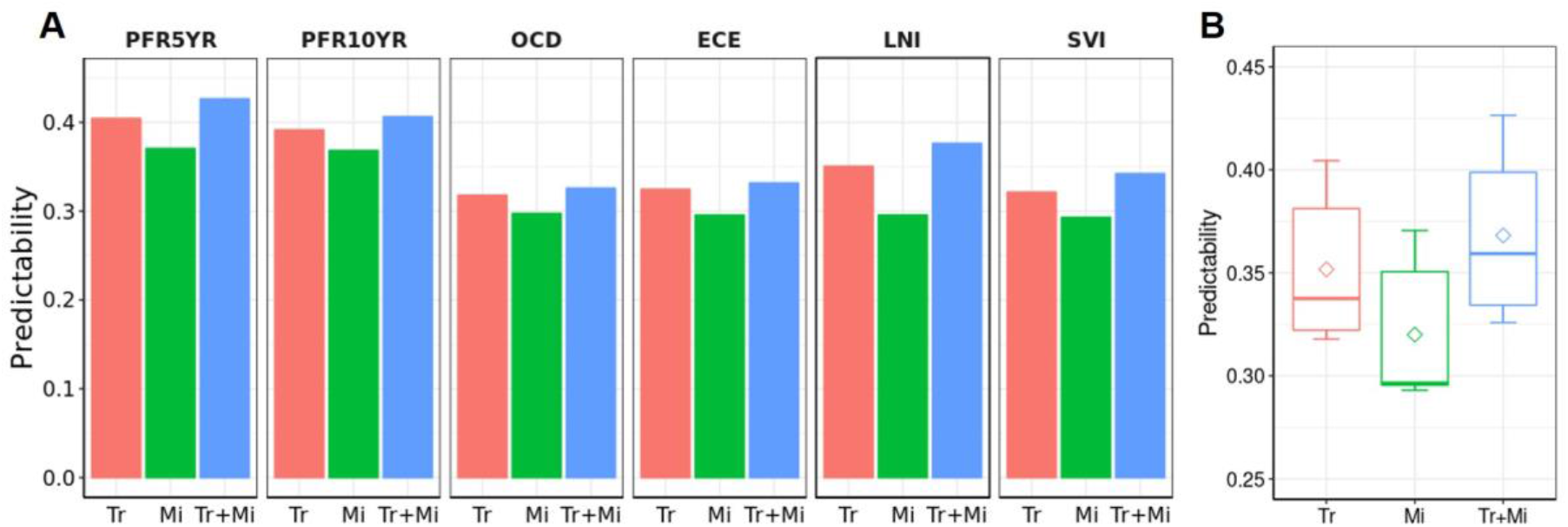
The performance of different expression panels in predicting the six nomogram-derived traits using BLUP-HAT. (**A**) Bar plot visualizing the predictability of each panel for predicting a trait. (**B**) Box plot visualizing the overall predictabilities of panels with different omics data across the six traits. (Tr: a panel of top genes with the highest predictability selected from the ranked gene list; Mi: a panel of top miRNAs with the highest predictability selected from the ranked miRNAs list; Tr+Mi: a combined panel of Tr and Mi. Genes/miRNAs in the Tr/ Mi panels for different traits are different)

### Development of multi-omics prognostic models by the SFS-BLUPH methodology

The predictive power and computational efficiency of the BLUP-HAT method have been demonstrated using six PCa outcome traits calculated by nomogram. We then leveraged this method to select a multi-omics signature for the prediction of RFS, the disease phenotype of interest. Patients with limited post-surgery follow-up data were eliminated from the initial 495 patients, leaving a total of 153 patients in this analysis, of which 95 underwent disease relapse or biochemical recurrence (BCR) within five years after prostatectomy. The outcome phenotypic value for a patient was defined as 1 if either this patient had not relapsed within five years or the time to first BCR was more than five years; otherwise, the outcome phenotypic value was calculated by dividing the time to first BCR by five, yielding a continuous score variable. Note that the greater the RFS score, the higher the probability of RFS (or the better the outcome). The newly defined outcome trait, which represented the probability of being RFS in five years (RFS5YR) after surgery, was most clinically relevant to disease prognosis.

In order to refine an optimal multi-omics signature for the prediction of RFS, we developed a novel stepwise forward selection strategy by leveraging the highly efficient BLUP-HAT method and the TCGA-PRAD multi-omics datasets. Similarly, we sorted all of the genes in descending order based on their absolute Pearson’s correlation coefficients with RFS. The initial BLUP-HAT model included the top two genes from the sorted list. In each following step, the next gene in the list was added to the current model for a calculation of the RFS predictability; this gene was retained if the addition of it increased the RFS predictability, otherwise, this gene was discarded. This selection process was repeated until all genes in the sorted list were sequentially tested, which yielded a refined 160-gene signature (GENE160) for predicting RFS. The same selection strategy was applied to the miRNA data to derive a refined 65-miRNA signature (MIR65) for predicting RFS.

In the TCGA-PRAD training set, three BLUP prognostic models (GENE160, MIR65, and GENE160+MIR65) were built using the selected genes and/or miRNAs for the prediction of the RFS scores. An RFS score was calculated for each patient *via* Leave-one-out cross validation (LOOCV), and the median value of these RFS scores was used to dichotomize the TCGA-PRAD cohort into a high-risk group (RFS scores less than the median value) and a low-risk group (RFS scores greater than the median value). The CoxPH regression analysis indicated that the scores calculated using all of the three signatures were significantly associated with RFS in the TCGA-PRAD training set (Figure 5A). The KM survival analysis showed that the patients in the low-risk group had significantly higher survival probability than those in the high-risk group (Figure 5B).

**Figure 5.**
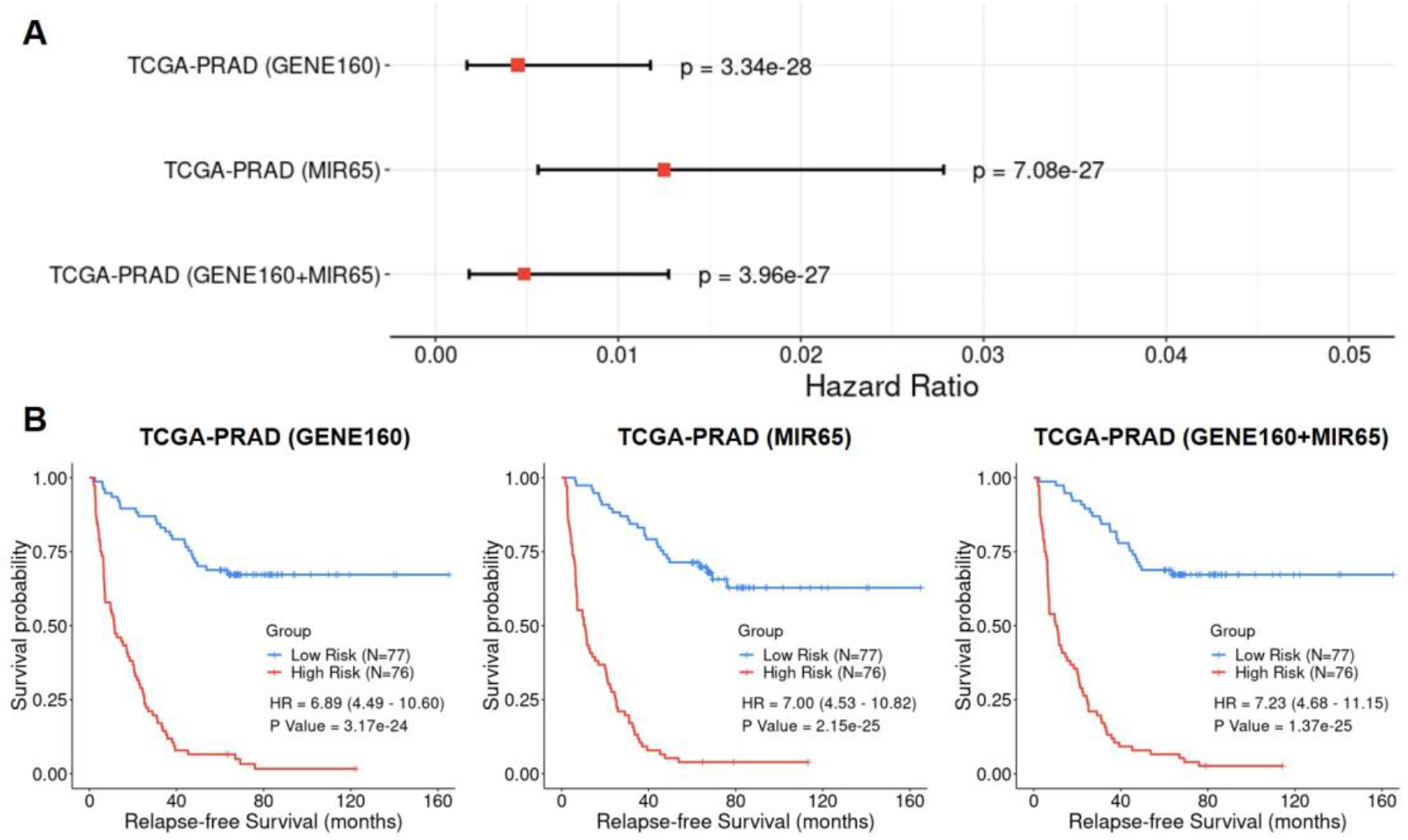
Cox Proportional-Hazards (CoxPH) and Kaplan-Meier (KM) survival analyses of relapse-free survival (RFS) using the GENE160, MIR65, and GENE160+MIR65 signatures in the TCGA-PRAD training dataset. (**A**) Forest plot visualizing the hazard ratio (HR), 95% confidence intervals, and p value of CoxPH survival analysis. (**B**) KM curves visualizing the survival probabilities over time for high and low risk groups classified based on the median predicted RFS scores in the cohort

We further validated the prognostic performance of the GENE160 and GENE160+MIR65 signatures using six independent cohorts. Note that these additional six datasets were not created using the same platform as the TCGA-PRAD data; thus, certain predictor variables of small number, either from 160 genes or from 65 miRNAs, were missing in some datasets (Supplementary Table S4). While validating the signatures and the methodology with each dataset, we only employed the available genes and/or miRNAs in a BLUP regression analysis. LOOCV was also used to calculate the RFS scores for the patients in each validation cohort. The CoxPH regression analysis and the KM analysis were then utilized to evaluate the association between the calculated RFS scores and the observed RFS outcomes. Although the RNAs were collected from different types of tissues (*i.e*., fresh frozen tumor tissue or FFPE) and the RNA abundance data were profiled using a variety of platforms (*i.e*., four different gene microarrays and RNAseq), the CoxPH regression analysis and the KM survival analyses indicated that the GENE160 signature alone was able to robustly predict RFS or differentiate high-risk patients from low-risk patients in these six datasets (Figure 6). Note that for the cohort of MSKCC2010, the CoxPH regression analysis rendered a significant result (p = 0.02), while the KM analysis only showed prognostic tendency (p = 0.15). Since the miRNA data is available for the MSKCC2010 dataset, we tested the multi-omics model with the integration of GENE160 and MIR65 signatures, which showed a significantly increased prognostic ability in this validation set. The p value for the CoxPH regression analysis has been improved from 0.02 (GENE160) to 5.76e-03 (GENE160+MIR65), while the result for the KM analysis became statistically significant (p = 0.019).

**Figure 6.**
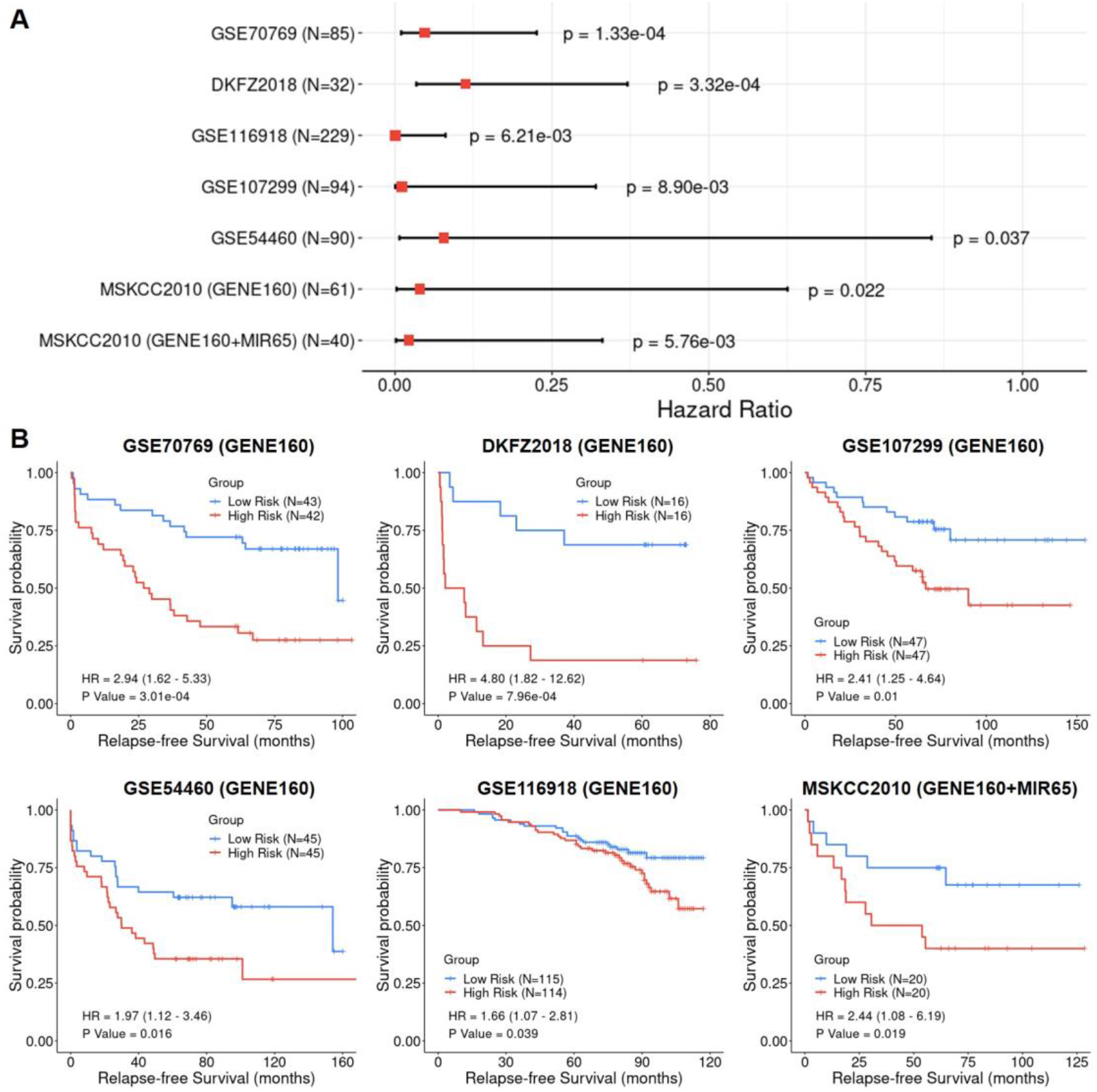
Cox Proportional-Hazards (CoxPH) and Kaplan-Meier (KM) survival analyses of relapse-free survival (RFS) using the GENE160 and GENE160+MIR65 panels in six independent validation datasets. (**A**) Forest plot visualizing the hazard ratio (HR), 95% confidence intervals, and p value of CoxPH survival analysis (**B**) KM curves visualizing the survival probabilities over time for high and low risk groups classified based on the median predicted RFS scores in each cohort.

## Discussion

Due to the cost of gene testing and the convenience of modeling, establishment of a prognostic test only using dozens of gene expression profiles has been the rule of thumb in the past decades. In our study, the predictabilities of three commercial panels of PCa prognosis were significantly higher than those of randomly selected gene sets, suggesting that the genes in these panels are indeed associated with disease progression. For example, Prolaris consists of 31 cell cycle progression (CCP) genes, many of which are functionally relevant to PCa recurrence [2]. Genes representing multiple biological pathways in Decipher are associated with PCa progression and have been reported to be differentially expressed throughout PCa progression [3]. The selected genes in Oncotype have also been verified to be related to PCa aggressiveness [4]. These several dozens of genes included in the commercial panels are no doubt biologically critical in PCa. However, these genes, even with major effects, may not be the best or complete set of predictors for PCa prognosis, which may be indicated by the results shown in Figure 3, *i.e*., all the sequentially evaluated models had better predictabilities than the three commercial gene panels. This may be ascribed to two major reasons: (1) due to the heterogeneity of PCa tumors, the major genes in one cohort may not necessarily be major players in another cohort, and (2) models with a large number of genes, including both major players and minor genes, may render a better prediction of outcomes than a panel with only ‘major genes’.

The rapid advancement in biotechnology has significantly reduced operational cost, allowing us to develop improved tests by including a large number of genes, a practice previously limited by economic constraints. However, conventional statistical methods cannot efficiently handle highly saturated models with *p » n, i.e*., the number of predictor variables is much larger than the number of observations. Robust GS models such as BLUP and Bayesian methods (*i.e*., BayesA, BayesB, and BayesC, etc.) have been proposed and applied to handle saturated linear regression models in plant and animal breeding. However, the computational advantages of these advanced methods have been rarely applied to cancer prognosis and warrant investigation. In this study, we took advantage of transdisciplinary expansion to adapt these powerful GS methodologies from agricultural sciences to human cancer research. The results indicated that BLUP outcompeted other rival methods in both predictive ability and computational efficiency. When many thousands of prediction models need to be compared, BLUP-HAT may further reduce the computational cost by avoiding lengthy CV.

The computationally efficient BLUP-HAT model was utilized to evaluate tens of thousands of models in regard to their performance in predicting clinical outcomes of PCa. The results from these comparisons demonstrated that, when compared with the currently used commercial panels with a limited number of genes, inclusion of many more genes with minor effects on the disease may collectively improve the overall RFS predictability. The BLUP-HAT model also enjoyed the easiness of combining multi-omics data into a single model, which allowed for a further improvement of the predictive ability.

We established a novel stepwise forward selection BLUP-HAT method to facilitate searching available multi-omics data for predictor variables with predictive potential. Using the TCGA data as a training set, we developed a 160-gene signature and a 65-miRNA signature for predicting the RFS of PCa. The GENE160 signature alone was successfully validated in all six independent cohorts, and the GENE160+MIR65 multi-omics signature showed significantly improved predictability compared with GENE160 signature in the only test set where miRNA data was available. Certain genes or miRNAs were missing in some validation sets because different platforms were used for generating these independent datasets. The RFS predictabilities in these validation analyses might have been increased if the missing genes/miRNAs were added back to the prognostic models. The validation was also successful when FFPE samples were analyzed (GSE116918 and GSE54460). These results indicated that the signatures and the methodology were robust even when the quality of RNA samples was relatively low, suggesting a great potential in clinical application. A limitation of the study is that the size of the training set (n = 153) and six validation sets (n < 100 in general) were small, which was quite different from studies of plants or animals. An improved prognostic model for an accurate prediction of RFS for PCa patients can be developed when data for large cohorts become available in the future.

In summary, we demonstrated that (1) a large number of disease-relevant genes render better prediction of PCa outcomes than a few dozen major genes, and (2) the combination of multi-omics predictor variables can further increase the predictability. We developed a novel SFS-BLUPH methodology which can efficiently search multi-omics data for predictor variables with prognostic potential. This method may be applied to any private database for the development of clinically useful tests for PCa prognosis. The new method may also be extendedly applied to different cancers or other types of human diseases.

## Biographical Note

Zhenyu Jia is an Associate Professor at University of California, Riverside, USA

Weide Zhong is a Full Professor at South China University of Technology, Guangzhou, China

Jianguo Zhu is a Urologist at Guizhou Provincial People’s Hospital, Guizhou, China

Ruidong Li, Han Qu, Le Zhang, Lei Yu and Meiyue Wang are PhD students at University of California, Riverside, USA

Shibo Wang and John M. Chater are Postdoctoral Fellows in Dr. Zhenyu Jia’s lab at University of California, Riverside, USA

Yanru Cui is an Associate Professor at Hebei Agricultural University, China

Yang Xu is an Assistant Professor at Yangzhou University, China

Julong Wei is a Postdoctoral Fellow at Wayne State University

Jianming Lu, Yuanfa Feng, and Rui Zhou are Ph.D. students in Dr. Weide Zhong’s lab at South China University of Technology, China

Yuhan Huang is a B.S. student at University of California, Los Angeles, USA

Renyuan Ma is a B.S. student at Bowdoin College, USA

## Key points

- We adopted genomic selection methods from the agricultural sciences and applied these to cancer research.
- We systematically evaluated the performance of six genomic selection methods using three omics data and their combinations in predicting prostate cancer outcomes, and found that the Best Linear Unbiased Prediction (BLUP) method outperformed the other models in terms of trait predictability and computational efficiency.
- With the more computationally efficient BLUP-HAT methodology, we demonstrated that (1) prediction models using expression data of a large number of genes selected from the transcriptome outperformed the clinically employed tests which only considered a small number of major genes, and (2) the integration of other omics data (*i.e*., miRNAs) in the model will further increase the predictability.
- We developed a novel stepwise forward selection BLUP-HAT (SFS-BLUPH) method to search multi-omics data for predictor variables to predict relapse-free survival of prostate cancer patients. The methodology has been successfully validated using six independent cohorts.

## Data Access

All the scripts used in this study, including data preprocessing, genomic selection model evaluation, implementation of BLUP-HAT method, development and validation of the SFS-BLUPH model, as well as data visualization are freely available at https://github.com/rli012/BLUPHAT.

## Acknowledgments

We would like to thank Professor Shizhong Xu at University of California, Riverside, for discussing this project with us and providing insightful comments on the study.

